# The Influence of Macrophages within the Tumor Microenvironment on Ovarian Cancer Growth and Response to Therapies

**DOI:** 10.1101/2025.01.29.635532

**Authors:** Parisa Nikeghbal, Danielle Burke, Dalet Armijo, Samuel Aldarondo-Quinones, Diane S. Lidke, Mara P. Steinkamp

## Abstract

While the majority of ovarian cancer (OC) patients respond to front-line carboplatin/paclitaxel chemotherapy and surgical debulking, nearly all patients will develop platinum-resistance and recur. Our study investigates how tumor-associated macrophages (TAMs) within the tumor microenvironment (TME) affect chemotherapy outcomes using OC patient-derived organoids and humanized patient-derived xenografts (huPDX). *In vitro* macrophage migration assays demonstrate the selective recruitment of M2 polarized macrophages to OC organoids. M2 macrophages, but not M1 macrophages, increase OC organoid viability and reduce their sensitivity to paclitaxel in co-culture assays. Furthermore, we identified BMS777607, a receptor tyrosine kinase inhibitor, that is capable of repolarizing M2 macrophages *in vitro* and reduces organoid viability by a macrophage-dependent mechanism. In a platinum-sensitive humanized patient-derived xenograft (huPDX) model, the presence of human immune cells increased between-mouse variability in response to paclitaxel with two of four mice demonstrating tumor regrowth after two weeks of treatment. A TAM-targeted CSF-1R inhibitor, BLZ945 significantly reduced the total number of human immune cells within the ascites fluid of huPDX3 mice, but did not reduce tumor burden. However BLZ945 in combination with paclitaxel reduced tumor burden with no sign of regrowth. Our study demonstrates that patient-derived OC organoids and huPDX, are useful for evaluating the immunomodulatory effects of therapies and the influence of TAMs on response. huPDX models could serve as a robust platform for preclinical testing of novel anti-cancer treatments, providing insights into the complex interplay between immune cells and cancer therapeutics.

## Introduction

Ovarian cancer (OC) is the most lethal gynecologic cancer in the United States, with 55% of patients diagnosed at a late stage when the cancer has already metastasized (1). Despite initial response to front-line treatment with surgery and platinum-taxane chemotherapy, 75% of patients will relapse with platinum-resistant disease (2). Five-year survival of distant metastatic OC patients is currently 31.4% (1), underscoring a need for innovative treatments for metastatic OC.

Recently, the use of immune checkpoint inhibitors (ICI) has revolutionized treatment for certain cancers, particularly in melanoma and endometrial cancer (3, 4). However, in OC, response rates to immune checkpoint inhibitors alone or in combination with chemotherapy have been poor, with overall response rates of 8% (KEYNOTE-100 trial) and 9.6% (JAVELIN Ovarian 100 trial) for the large OC clinical trials (5). One major factor limiting response to ICIs in OC may be the immunosuppressive tumor microenvironment (TME).

Macrophages become polarized based on signals from their environment, and can adopt an anti-tumor (M1-like) or pro-tumor (M2-like) phenotype (6, 7). The ratio of M1 to M2 tumor-associated macrophages (TAMs) has been correlated with clinical outcomes across various solid tumors, including OC, where a higher ratio is associated with improved five year survival (8, 9). However, the OC TME appears to be populated by predominantly tumor-supporting TAMs. OC cells produce cytokines that recruit tumor-supporting (TAMs) to the TME. Pro-tumor TAMs make up 30-50% of the cells in OC malignant ascites (10, 11). TAMs in turn promote the recruitment of other immunosuppressive cells such as regulatory T cells and myeloid-derived suppressor cells (MDSCs), which further inhibit effective T cell responses (12). TAMs secrete cytokines that promote Th2 cell activity and directly suppress T cell function through PD-L1 and B7-H4 (12–15). TAMs also secrete growth factors that promote OC growth. Co-culturing OC cell lines with THP-1 derived macrophages has been shown to increases proliferation and migration by activating EGFR signaling (16–18).

There is an association between TAMs and chemotherapy response. Paclitaxel, one of the standard chemotherapy agents for front-line OC treatment, has been shown to alter the TME by promoting macrophage recruitment and polarization. In the MMTV-PyMT transgenic mouse model of breast cancer, paclitaxel treatment was associated with increased CSF1 expression and enhanced macrophage recruitment (19). Conversely, TAMs appear to promote OC resistance to carboplatin. Co-culturing OVCAR3 ovarian cancer cells with CD68+ macrophages in a hetero-spheroid model demonstrated enhanced resistance to carboplatin treatment (20).

In this study, we take a close look at human TAM/OC cell reciprocal interactions in patient-derived models of OC. We use an *in vitro* macrophage migration assay to examine between-patient variability in macrophage recruitment. We also use *in vitro* 3D co-culture systems that more accurately represent the TME (21). In addition, we used *in vivo* humanized PDX (huPDX) to investigate the effect of macrophages on OC chemotherapy response. By examining both *in vitro* dynamics and *in vivo* interactions, we have demonstrated that M2 polarized TAMs are selectively recruited by OC cells and that these pro-tumor TAMs can significantly reduce the effectiveness of paclitaxel. We show that a CSF1R inhibitor, BLZ945, in combination with paclitaxel limits paclitaxel resistance. We also identify a small molecule inhibitor, BMS777607, that alters macrophage polarization and inhibits OC organoid viability using a macrophage-dependent mechanism. These findings underscore the potential of targeting the immune environment, particularly TAMs, to improve outcomes in OC treatment.

## Material and Methods

### Cell lines and treatment

The THP-1 human monocyte cell line, derived from an acute monocytic leukemia patient, was obtained from ATCC. The OC cell line OVCAR8 was sourced from the NCI Cancer Repository. Both cell lines were authenticated using Short Tandem Repeat (STR) analysis provided by ATCC and were cultured in RPMI 1640 (Invitrogen) supplemented with 10% fetal bovine serum (FBS), 1% L-glutamine, and 1% penicillin-streptomycin, maintained at 37°C. Human monocytes were isolated from cryopreserved PBMCs (AllCells) using the Dynabeads Untouched Human Monocytes Kit (Thermo Fisher).

### Differentiation and Macrophage Polarization

THP-1 cells were seeded in suspension flasks at a density of 3 × 10^5^ cells/ml in RPMI media supplemented with 10% FBS, 1% L-glutamine, 1% Pen/Strep, and 50 uM ß-mercaptoethanol as specified by ATCC. For macrophage differentiation, 5 × 10^6^ THP-1 cells were seeded into a T25 flask supplemented with 150 nM phorbol 12-myristate 13-acetate (PMA, MilliporeSigma #1839) for 24 hours, followed by a 24-hour incubation in PMA-free medium supplemented with 50 uM ß-mercaptoethanol (22). Polarization was initiated on the third day. M1 macrophages were induced with 20 ng/mL IFN-γ (PeproTech Catalog # 300-02) and 250 ng/ml lipopolysaccharide (LPS, MilliporeSigma Catalog # L4524). M2 macrophages were induced with 30 ng/ml interleukin-4 (IL-4, PeproTech #200-04), for 48 hours. THP-1 polarization was routinely assessed by harvesting and labeling macrophages with the M1 marker CD80 and the M2 marker, CD206 (**Supplemental Fig. 1 A-B**)

PBMC-derived monocytes were differentiated using ImmunoCult-SF Macrophage Differentiation Media (STEMCELL Technologies) supplemented with 50 ng/ml M-CSF (Thermo Fisher Catalog #300-25-10UG). Polarization was initiated on the fourth day. M1 macrophages were induced with 50 ng/ml IFN-γ and 10 ng/ml LPS, while M2 macrophages were induced with 10 ng/ml IL-4, for 48 hours. Polarization of PBMC-derived macrophages was assessed by flow analysis with CD80 and CD206 as with THP-1-derived macrophages (**Supplemental Fig. 1 C-D**)

### Patient-Derived Xenograft Organoid (PDXO) Formation

Organoid seeding followed protocols based on those developed in the Leslie (University of New Mexico) and Thiel (University of Iowa) labs (23, 24). Briefly, fresh or cryopreserved PDX tumors were finely minced and then digested in digestion buffer (2 U/mL dispase (Millipore Sigma D4693-1G), 1 mg/mL collagenase P (Millipore Sigma 11213857001), and 2 µL/ml DNAse in human wash buffer) for 30 minutes at 37°C on a tube rotator. The digested tissue was strained through a 40 µm strainer into a 15 mL tube, then washed twice with human wash buffer (Phenol red-free DMEM/F12 (GIBCO, #21041025) supplemented with 1 M HEPES (pH 7.2-7.5), 100X GlutaMAX, 50 mg/mL Primocin, and 10% BSA), with centrifugation at 500xg for 10 minutes after each wash. For organoid seeding, 50,000 single cells were embedded in 100% ultimatrix RGF Basement Membrane Extract (#BME001-10, Cultrex) in 50 µL domes in a 24-well plate. After 20 minutes of incubation, 450 µL of organoid media was added. The media is composed of Advanced DMEM/F12 (Gibco, 12634028) supplemented with 1X GlutaMAX, 10 mM HEPES, and Pen/Strep and is enriched with 100 ng/ml recombinant human Noggin (R&D Systems Catalog #6057-NG-025), 250 ng/ml R-spondin-1 (R&D Systems Catalog # 4645-RS-025), 100 ng/ml human FGF-10 (ThermoFisher Scientific Catalog #100-26-1MG), 37.5 ng/ml Heregulin beta-1 (ThermoFisher Scientific Catalog #100-03-1MG), 50 ng/ml EGF (ThermoFisher Scientific, Catalog # AF-100-15-1MG), B27 supplement (1:50, ThermoFisher Scientific, Catalog #17504044), 1.25 mM N-Acetylcysteine (Cayman Chemical, Catalog # 616-91-1), 100 nM beta-estradiol (Millipore Sigma, Catalog # 50-28-2), 2% Primocin (InvivoGen, Catalog # ant-pm-1), 5 mM Nicotinamide (Millipore Sigma, Catalog #N0636-500G), 5 µM A83-01 (STEMCELL Technologies, Catalog # 72024), 10 µM Y-27632 (Sigma Aldrich, Catalog # Y0503-1MG),10 µM Forskolin (Bio-Techne, Catalog # 1099), and 250 µg/ml Hydrocortisone (STEMCELL Technologies, Catalog # 74144). Organoid formation typically occurs between 6 to 12 days, depending on the cell source.

### *In vitro* 3D Monocyte/Macrophage Migration Assay

OVCAR8 spheroids or PDX organoids were harvested and seeded in a layer of VitroGel Organoid 3 synthetic matrix (The Well Bioscience), a hydrogel optimized for organoid cultures, at a 1:1.2 ratio (100 µl complete RPMI media:120 µl VitroGel) in an 8-well chamber. 100 µL of complete RPMI medium was added to each well. After 24 hours, 100,000 PBMC-derived or THP-1 derived monocytes, M1, or M2 macrophages following previously described protocols for differentiation and polarization, were added on top of the matrix. After 3 days of co-culture, organoids were collected and fixed in 4% PFA. Macrophages were labeled with anti-human CD45-Alexa532 antibody and nuclei were stained with DAPI for whole-organoid imaging using a Zeiss LSM 800 confocal microscope. The number of infiltrating macrophages/organoid was quantified from organoid z-stack images and normalized to the organoid area (in µm^2^) of each image. Normalization was required due to the larger size of organoids with infiltrating M2 macrophages and the variability in organoid sizes among different PDX.

### Co-culture of TAMs and OC cells

#### Monolayer Co-culture

For monolayer cultures, 5,000 OVCAR8 cells/well were seeded into a 96-well flat-bottom plate and allowed to adhere for 6 hours. THP-1 derived macrophages were fluorescently labeled for 30 minutes with Cell Tracker Green (Thermo Fisher) at a 1:20 dilution and rinsed in PBS. 500 macrophages were added per well. 24 hours later, co-cultures were treated with PTX at 1 nM, 5 nM, 10 nM, or 50 nM and incubated for 3 days in the Incucyte S3 with images captured every 12 hours. Change in confluency was analyzed longitudinally using the IncuCyte S3 imaging system (Sartorius). Difference in confluency with or without treatment and with or without macrophages was assessed after 72 hours of treatment. All assays tested 4 wells/condition as technical replicates. Three independent drug response studies were run as biological replicates.

#### Spheroid Co-culture

THP-1-derived or PBMC-derived macrophages and OVCAR8 cells were mixed at a ratio of 1:10 macrophages to OVCAR8 cells and seeded into 96-well cell-repellent U-bottom plates (Greiner Bio-One). Spheroids formed over 24 hours. Spheroids were then treated with paclitaxel at concentrations of 1 nM, 5 nM, 10 nM, or 50 nM for 3 days. At the endpoint, cell viability was assessed using the CellTiter-Glo® 3D Cell Viability Assay (Promega, Catalog # G9682), which is specifically designed for analysis of three-dimensional cell culture models. All assays tested 4 wells/condition as technical replicates. Three independent drug response studies were run as biological replicates.

#### Organoid Co-culture

Organoid co-cultures were established in 96-well plates, seeding 20-30 organoids per well (24). PDXOs were prepared in 24-well plates, harvested, washed, and resuspended in cold media mixed with Ultimatrix at a 1:1 ratio, then co-cultured with 2500 macrophages per well in 5 µL domes. After solidification, 200 µL of co-culture media (comprising 50% organoid media without hydrocortisone and forskolin and 50% ImmunoCult macrophage media with 50 ng/ml M-CSF and 1:50 Insulin-Transferrin-Selenium-X (Gibco #1748235)) was added. Treatment with chemotherapy agents (Carboplatin, Paclitaxel, or their combination) and BMS777607 at specified concentrations followed the next day. 0.1% DMSO, diluted in media, served as the vehicle control. Cell viability was assessed 72 hours post-treatment using the CellTiter-Glo® 3D Cell Viability Assay, which measures ATP as an indicator of metabolic activity. An equal volume of CellTiter-Glo® 3D reagent was added to each well, and after vigorous shaking, plates were allowed to stabilize at room temperature for 25 minutes before measuring luminescence with a BioTek Synery Neo2 Multi-Mode Plate Reader. Each condition was tested with six or more technical replicates, and two independent experiments were conducted per organoid culture as biological replicates.

### Macrophage Repolarization Assay

THP-1 monocytes were seeded in T25 flasks at a density of 5 × 10^6 cells per flask. Following established protocol, they were polarized to M1 and M2 phenotypes. Subsequently, these cells were then treated with various concentrations of repolarizing drugs for 72 hours to assess potential phenotype shifts. After treatment, macrophages were detached using Accutase (STEMCELL Technologies) and analyzed by flow cytometry to evaluate repolarization efficacy.

### Establishing PDX Models from Malignant Ascites

All mouse procedures were approved by the UNM Animal Care and Use Committee, in accordance with NIH guidelines for the Care and Use of Experimental Animals. PDX models were established from malignant ascites collected under IRB-approved protocol INST1509 from the University of New Mexico as detailed in Steinkamp et al., 2023 (25). Briefly, isolated primary cancer spheroids were injected intraperitoneally into NSG mice (RRID: IMSR_JAX:005557) to establish orthotopic PDX models of disseminated OC as approved under IACUC protocol #21-20114-HSC. Authentication of PDX lines was confirmed through STR profile analysis, comparing PDX alleles with those of primary patient samples to ensure derivation from the patient tumors.

### Humanized NBSGW Mice

Humanized mice were established by the UNMCCC Animal Models Shared Resource under IACUC-approved protocol 22-201300-HSC, as described previously (25). In brief, NBSGW mice (RRID:IMSR_JAX:026622) were engrafted with cord blood-derived human CD34+ hematopoietic stem cells (HSC). Cryopreserved HSCs were thawed, and incubated in Stem Line II medium (Sigma-Aldrich) with human recombinant SCF (STEMCELL Technologies) for 6 hours, and then resuspended in PBS. Three-week-old NBSGW females received 7.5 × 10^4^ CD34+ HSC cells via retro-orbital injection. Humanization was confirmed by flow cytometry, assessing the percentage of human CD45+ cells versus mouse CD45+ cells in the peripheral blood 16 weeks post-engraftment. huNBSGW mice were divided into six treatment groups. Mice with high and low percentage humanization based on PBMC analysis at 16 weeks post-engraftment were distributed equally across groups (average humanization=51.5% with a range from 20-86%). Humanization for all mice is plotted in **Supplemental Fig 2**.

### Non-Humanized PDX and Humanized PDX Treatment and Tumor Monitoring

Humanized and non-humanized NBSGW mice were injected intraperitoneally (IP) with 5 × 10^6^ PDX3-Luc2 as spheroids. After two weeks of tumor growth, pre-treatment tumor burden was quantified by bioluminescent imaging on the IVIS Spectrum *in vivo* imaging system. Non-humanized PDX (non-huPDX) mice (n=6/treatment group) were treated with carboplatin (CBDCA, 12 mg/kg, twice weekly), low-dose paclitaxel (PTX, 6 mg/kg, 5 days a week), and the combination CBDCA plus PTX. The control group was treated with DMSO diluted in PBS. HuPDX mice (n=4 or 5/treatment group) were treated with the same regimen of CBDCA, PTX, or CBDCA plus PTX and two additional groups treated with the CSF1R inhibitor BLZ945 (sotuletinib)(160 mg/kg, 5 days a week) or BLZ945 plus low-dose paclitaxel to test the effect of a macrophage-targeting therapy on response in the huPDX model.

Tumor burden was monitored weekly by bioluminescent imaging on the IVIS Spectrum *in vivo* imaging system (Perkin Elmer). D-Luciferin was administered IP and mice were imaged 10 minutes post-injection under isoflurane anesthesia. Imaging parameters were standardized to ensure consistent signal acquisition. Response to treatment was measured as a change in bioluminescent signal intensity from pre-treatment tumor burden after two or three weeks of treatment. Non-humanized PDX3 mice were treated for three weeks and then sacrificed when control mice had high tumor burden and abdominal swelling due to ascites fluid. For humanized PDX3, we chose to sacrifice mice after two weeks of treatment to limit attrition due to the development of anemia and wasting that can occur in humanized PDX models. At this time point, we were able to perform spectral flow analysis on fresh ascites samples from all groups.

### Spectral Flow Cytometry Analysis of huPDX3 Ascites Fluid

Spectral flow cytometry was conducted to analyze immune cells isolated from the ascites fluid of OC patients at the time of surgery and huPDX and non-huPDX mice at the experimental end point. A 15-color panel was used to profile major immune cell subsets. This panel included 13 human-specific markers targeting immune cell surface proteins and a viability stain (**Table S1**, all antibodies from BioLegend, San Diego, CA). For humanized PDX samples, a human-specific CD45 antibody and a mouse-specific anti-CD45 antibody were used to select human immune cells and gate out mouse immune cells. Fluorophores were distributed across 4 lasers to limit overlap in co-expressing cells. Similarity indices for each fluorochrome was calculated with the Cytek Full Spectrum Viewer (Cytek) and the Complexity Index was determined to be 7.6. Ascites fluid was collected from patients at the time of surgery or mice at necropsy, and live cells were centrifuged at 500 x g for 10 minutes at 4°C and red blood cells were lysed with ammonium chloride (STEMCELL Technologies, Vancouver, BC). Cells were rinsed with RPMI media, filtered through cell strainers to remove cancer spheroids and to obtain a single cell suspension, and counted on the Countess II (Thermo Fisher Scientific). Cells were resuspended in 1 mL FACS Buffer (1% FBS in PBS). Human Fc receptors were blocked with 10 μg/mL Human BD Fc Block (BD Pharmingen) for 10 minutes at room temperature. The 14-antibody cocktail was diluted in FACS buffer supplemented with Brilliant Stain Buffer (BD Horizon) and cells were stained for 1 hour on ice, protected from light. After removing the unbound antibodies, cells were stained with Zombie NIR Fixable Viability Kit (BioLegend) at 1:10,000 dilution for 15 minutes at room temperature to label dead cells. A live-dead single stain control was prepared by heating unstained cells to 65°C for 10 minutes to kill cells, followed by staining with Zombie NIR. All samples were washed in FACS buffer and filtered through 5 ml round bottom tubes with cell-strainer cap (352235, Falcon). Phenotyping was performed using the Cytek® Aurora Full Spectrum Flow Cytometer, and then analyzed using SpectroFlo® Flow Cytometry Software v3.0 (Cytex Biosciences, Fremont, CA, USA) and FlowJo (10.6) (Becton, Dickinson and Company; Franklin Lakes, NJ). Ultra Comp eBeads (01-2222-42, Invitrogen) were used for single stain controls. An autofluorescence signature identified in unstained cells was designated as an additional fluorescent tag in unmixing. Manual gating removed doublets, debris, and dead cells. The human CD45+ subset was identified for each mouse, randomly down-sampled to 10,853 cells to obtain equal cell numbers across samples, and concatenated into one file (n=20) for further analysis. Human CD45+ immune cells from all treatment groups were further analyzed by T-distributed stochastic neighbor embedding (tSNE) to cluster immune cell subpopulations. Due to early removal of mice due to wasting/anemia, analysis included control mice (n=4) and treated mice: paclitaxel (n=4), carboplatin (n=2), paclitaxel + carboplatin (n=4), BLZ945 (n=2) and paclitaxel + BLZ945 (n=4).

### Statistical Analyses

Between group analyses were performed using GraphPad Prism Version 10.4.1. One way ANOVA was used to determine whether differences between groups was significant (p<0.05).

### Data Availability

The data generated in this study are available within the article and its Supplementary Data.

## Results

### Macrophage Recruitment by OC Organoids

OC cells produce a variety of chemokines such as macrophage colony-stimulating factor (M-CSF) and CC-chemokine ligand 2 (CCL2/MCP-1) that can influence the recruitment and distribution of macrophages within the TME (12, 25). These factors can also promote macrophage polarization towards a pro-tumor phenotype, supporting an immunosuppressive tumor microenvironment (26, 27). Our previous paper identified PDX-specific cytokine signatures in three OC PDX models (25). To understand how differential production of secreted cytokines influences macrophage recruitment to the TME, we developed an *in vitro* 3D macrophage migration assay. PDX-derived organoids were embedded in vitrogel, a synthetic matrix, and PBMC-derived monocytes or macrophages: M1 (pro-inflammatory, anti-tumor) or M2 (anti-inflammatory, pro-tumor), were seeded on the matrix surface and allowed to migrate to the organoids (**Fig 1A**). After 3 days, fixed organoids were labeled by IF with an anti-CD45-Alexa532 antibody to detect monocytes or macrophages within the organoids (**Fig 1B**). Six models, the OVCAR8 ovarian cancer cell line and five PDX-derived organoids, were tested over two experiments using separate macrophage donors (**Fig 1C**, D). PDXO9, tested in both experiments, demonstrated consistent recruitment, reinforcing the assay’s consistency (**Fig 1C**, D). In four out of six models, organoids incubated with M2 macrophages were larger than those incubated with M1 macrophages or monocytes, (**Fig 1C**). To account for the difference in organoid size, the number of monocytes or macrophages within each organoid was normalized to the maximum estimated area of that organoid. We were able to identify an infiltrating monocytes and macrophages in PDXOs, in agreement with a recent microfluidics study that observed M0 macrophages infiltrating into a collagen gel seeded with single-cell OC cells (28). Uniquely, we found that across all models, M2 macrophages infiltrated organoids at much higher numbers compared to monocytes or M1 macrophages (**Fig 1D**). The total number of infiltrating M2 macrophages also varied among the models with PDXO 3 having the highest number of infiltrating M2 macrophages and the OVCAR8 cell line and PDXO 9 having the least. Interestingly, the number of infiltrating M2 macrophages correlated with the level of M-CSF detected in the patients’ primary ascites fluid as measured by ELISA (**Fig 1E**). Organoids derived from patients with elevated M-CSF, Patients 3, 18 and 20, showed a corresponding higher recruitment of M2 macrophages (**Fig 1D**, E). Conversely, organoids derived from patients with lower M-CSF, Patients 2 and 9, recruited fewer M2 macrophages (**Fig 1D**, E). This correlation suggests that M-CSF may be a key driver in the preferential recruitment of M2 macrophages to the TME. Other chemokines involved in macrophage and/or monocyte recruitment (MCP-1, IP-10 and MIG) did not show a correlation with monocyte/macrophage infiltration (**Fig 1E**).

**Figure 1.**
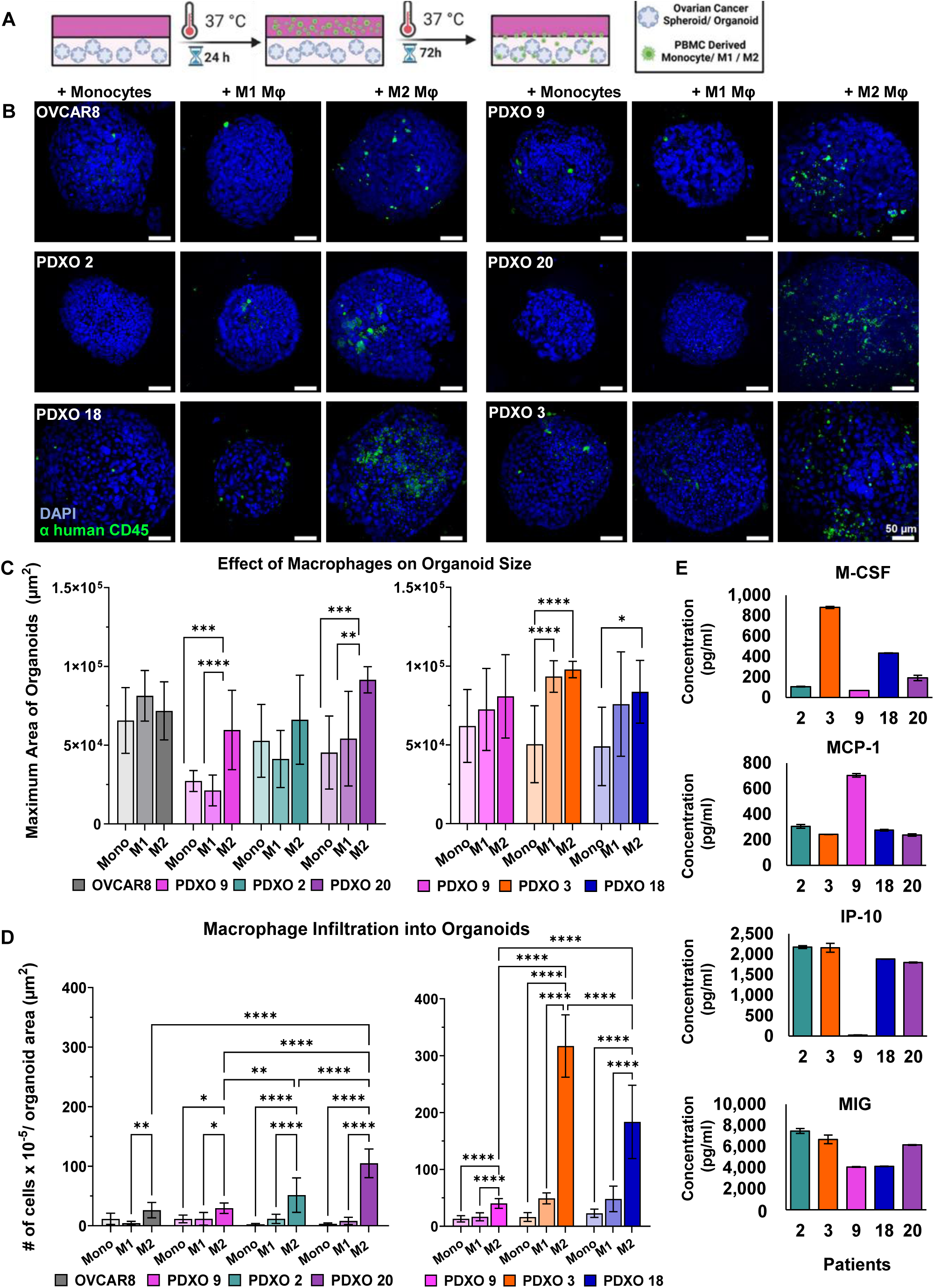
Macrophage Recruitment by OC Organoids. **A.** Schematic of the macrophage migration assay: OVCAR8 spheroids or PDXOs were seeded in a layer of Vitrogel Organoid 3 matrix. PBMC-derived monocytes (Mono), M1 or M2 macrophages were added on top of the gel surface. After 72 hrs., organoids were fixed and imaged. **B**. Representative confocal images of fixed organoids. (Blue: DAPI, Green: CD45+ monocytes/macrophages, Scale bar = 50 μM). **C.** The average size of organoids following the migration assay (n=10 per condition). **D.** The total number of CD45+ cells/organoid was normalized to the maximum organoid area. Significant differences are shown (lines, one-way ANOVA, ** p<0.01, *** p<0.001, **** p<0.0001). **E**. Cytokine concentrations in matched patient ascites fluid based on ELISA. Values are the average of two technical duplicates.

These findings indicate that OC organoids preferentially recruit M2 macrophages, rather than monocytes or M1 macrophages. The extent of this recruitment may be primarily modulated by the concentration of M-CSF present in the TME. Thus, a patient’s cytokine signature likely influences macrophage recruitment, thereby affecting the tumor’s progression and response to therapy.

### Impact of Macrophage Polarization on OVCAR8 Response to Palitaxel Therapy

To investigate how human macrophages influence chemotherapy response in OC and to determine whether the 3D microenvironment influences this response, platinum-resistant OVCAR8 cells were co-cultured with M1 or M2 macrophages as 2D monolayer (**Fig 2A, B**) or 3D spheroid (**Fig 2E**) cultures and then treated with the standard chemotherapy drug paclitaxel.

**Figure 2.**
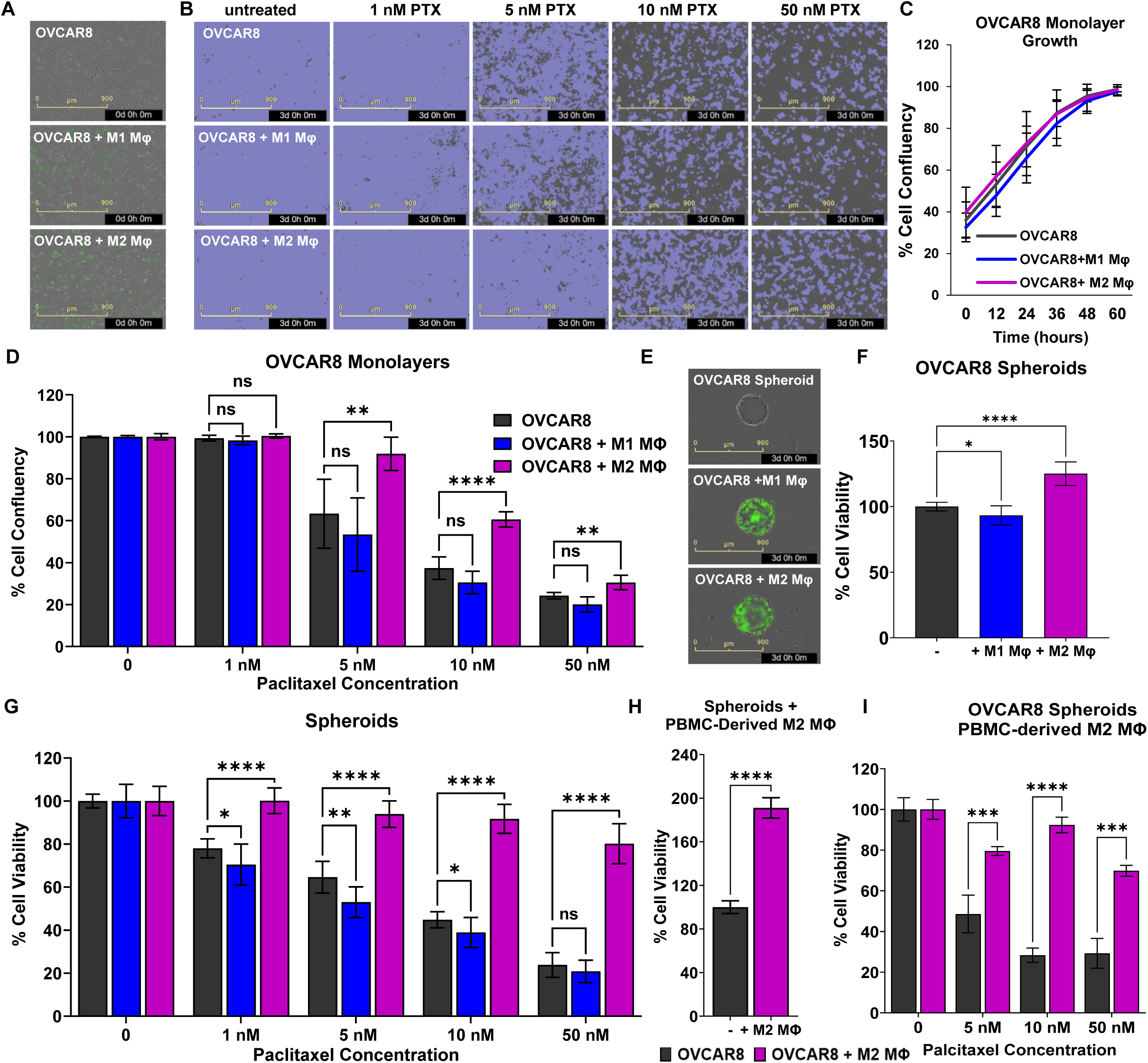
Impact of Macrophage Polarization on OVCAR8 Response to Paclitaxel. OVCAR8 cells were co-cultured with M1 and M2 macrophages derived from THP-1 cells **(A-G)** or PBMCs **(H-I) A.** Overlaid phase contrast and fluorescent images of untreated OVCAR8 cells +/- THP-1-derived M1 or M2 macrophages (Mφ, green) as monolayers 24 hours after seeding at a 10:1 cancer cell to macrophage ratio. Scale bar =900µm **B.** Representative images of OVCAR8 cell confluence (purple) +/- M1 or M2 THP-1-derived macrophages after 3 days of treatment with 0,1, 5, 10, or 50 nM PTX. A confluence mask was overlaid on a phase contrast image of the cells. Scale bar =900µm **C.** Change in cell confluency of untreated OVCAR8 cells seeded as a monolayer with or without M1 or M2 macrophages. **D**. Cell confluency normalized to untreated controls (100%) in each group (no macrophages, M1, or M2 macrophages) after 3 days of paclitaxel treatment**. E.** Overlaid phase contrast plus fluorescent images after 3 days of co-culturing untreated OVCAR8 spheroids +/- THP-1-derived M1 or M2 (green) macrophages seeded at a 10:1 ratio in u-bottom cell repellent plates. Scale bar =900µm **F.** Effect of M1 or M2 macrophages on the viability of untreated spheroids after 3 days of co-culturing. **G.** OVCAR8 spheroid viability normalized to untreated control (100%) in each group after 3 days of paclitaxel treatment. **H.** Difference in spheroid viability of untreated OVCAR8 +/- PBMC-derived M2 macrophages after 3 days. **I.** Spheroid viability in OVCAR8 +/- PBMC-derived M2 macrophages normalized to untreated control (100%) in each group and treated for 3 days with paclitaxel. (* p<0.05, ** p< 0.01, *** p< 0.001, **** p<0.0001, using a one-way ANOVA). For all assays (D-I), 4 wells were tested/condition (technical replicates) and three independent experiments (biological replicates) were averaged

OVCAR8 cells were co-cultured as monolayers with or without CellTracker Green-labeled THP-1 derived macrophages in a 10:1 ratio of cancer cells to macrophages. OVCAR8 alone, or co-cultured with M1 or M2 macrophages, exhibited similar cell confluency 24 hours after seeding (**Fig 2A**). Cells were then treated with a range of concentrations of paclitaxel for 3 days and evaluated for cell confluency using the Incucyte S3 imaging system (**Fig 2B**). No significant difference in cell confluency was observed in untreated OVCAR8 alone, or co-cultured with M1 or M2 macrophages over time (**Fig 2B, C**). Paclitaxel treatment reduced the growth of OVCAR8 cells in a concentration-dependent manner, confirming OVCAR8 sensitivity. OVCAR8 cells co-cultured with M2 macrophages exhibited higher confluency when treated with 5 nM, 10 nM and 50 nM paclitaxel, compared to OVCAR8 cells grown alone or with M1 macrophages (**Fig 2D**). Thus M2 macrophages reduced OVCAR8 sensitivity to paclitaxel. However, higher concentrations of paclitaxel still reduced cell confluency in OVCAR8 cells co-cultured with M2 macrophages.

To determine the effect of the 3D morphology on macrophage-dependent paclitaxel resistance, chemotherapy response was further assessed in spheroid cultures, which model free floating OC spheroids present in patient ascites (**Fig 2E**). OVCAR8 cells were seeded into cell-repellent U bottom plates with or without CellTracker Green-labeled THP-1-derived macrophages. Condensed spheroids formed by 24 hours. Spheroids were administered varying concentrations of paclitaxel and cell viability was assessed after 72 hours of treatment. Co-culturing OVCAR8 spheroids with M2 macrophages significantly improved cell viability by 1.25-fold, while M1 macrophages slightly reduced viability (**Fig 2F**). Because macrophages altered growth in untreated spheroids, response to paclitaxel was normalized to the untreated control for each condition: no macrophages, M1 or M2 macrophages. OVCAR8 spheroids co-cultured with M2 macrophages retained 80% or greater viability across all tested paclitaxel concentrations, compared to OVCAR8 spheroids with no macrophages. The effect of M2 macrophage co-culture in paclitaxel sensitivity was more pronounced in the spheroid studies than in the monolayer studies (**Fig 2D, G**). When treated with the highest concentration of paclitaxel, average viability was reduced by only 20% in spheroids incubated with M2 macrophages, while in monolayer co-cultures, average confluency was reduced by 70%. M1 macrophages also had a significant negative effect on spheroid viability, which was not seen in monolayer studies (**Fig 2G**).

To ensure that these effects were not specific to the THP-1-derived macrophages, co-culture studies were repeated with human PBMC-derived M2 macrophages. Co-culture with these M2 macrophages increased untreated OVCAR8 cell viability by two-fold (**Fig 2H**) and significantly reduced OVCAR8 sensitivity to paclitaxel for all concentrations, maintaining greater than 70% viability. This highlights the pro-tumorigenic activity of human M2 macrophages in promoting cell survival with paclitaxel treatment (**Fig 2I**). This *in vitro* data suggests that targeting tumor associated macrophages could potentially improve therapeutic outcomes especially in patients that receive second line paclitaxel monotherapy. The differential impact of macrophage phenotypes on cancer cell viability underscores the complexity of the TME and the need for tailored therapeutic strategies that consider the complex interplay between cancer cells and immune cells, particularly TAMs whose recruitment is likely patient-dependent.

### Impact of Macrophage Polarization on Response to Treatment in PDXO Models

We next wanted to test whether M2 macrophages can influence chemotherapy response in more clinically relevant patient-derived models that each have unique genetic signatures. Organoids were established from three distinct PDX, two PDX drived from platinum-sensitive patients (PDX3, PDX18) and one from a platinum-resistant patient (PDX9). For baseline measurements, the initial viability at Day 0 of organoids seeded alone or in the presence of THP-1-derived macrophages was determined (**Supplemental Fig. 3**). The difference in viability between organoids alone and organoids seeded with macrophages identified the component of the whole-well viability measurement that represents the metabolic activity of the macrophages themselves. The addition of M1 or M2 macrophages to PDXO3, PDXO9, or PDXO18 organoid cultures increased the initial viability measurement by an average of 14.2% and 14.9% respectively, when compared to wells seeded with organoids alone. After three days of co-culture, M2 macrophages significantly enhanced the viability of organoids across all three PDXO models, increasing viability by an average of 28.2% (PDXO3), 42.7% (PDXO18), and 27.8% (PDXO9) compared to organoids alone. On the other hand, M1 macrophages did not increase organoid viability in any co-cultures and instead significantly reduced viability by an average of 15.5% in PDXO3 (**Fig 3A**).

**Figure 3.**
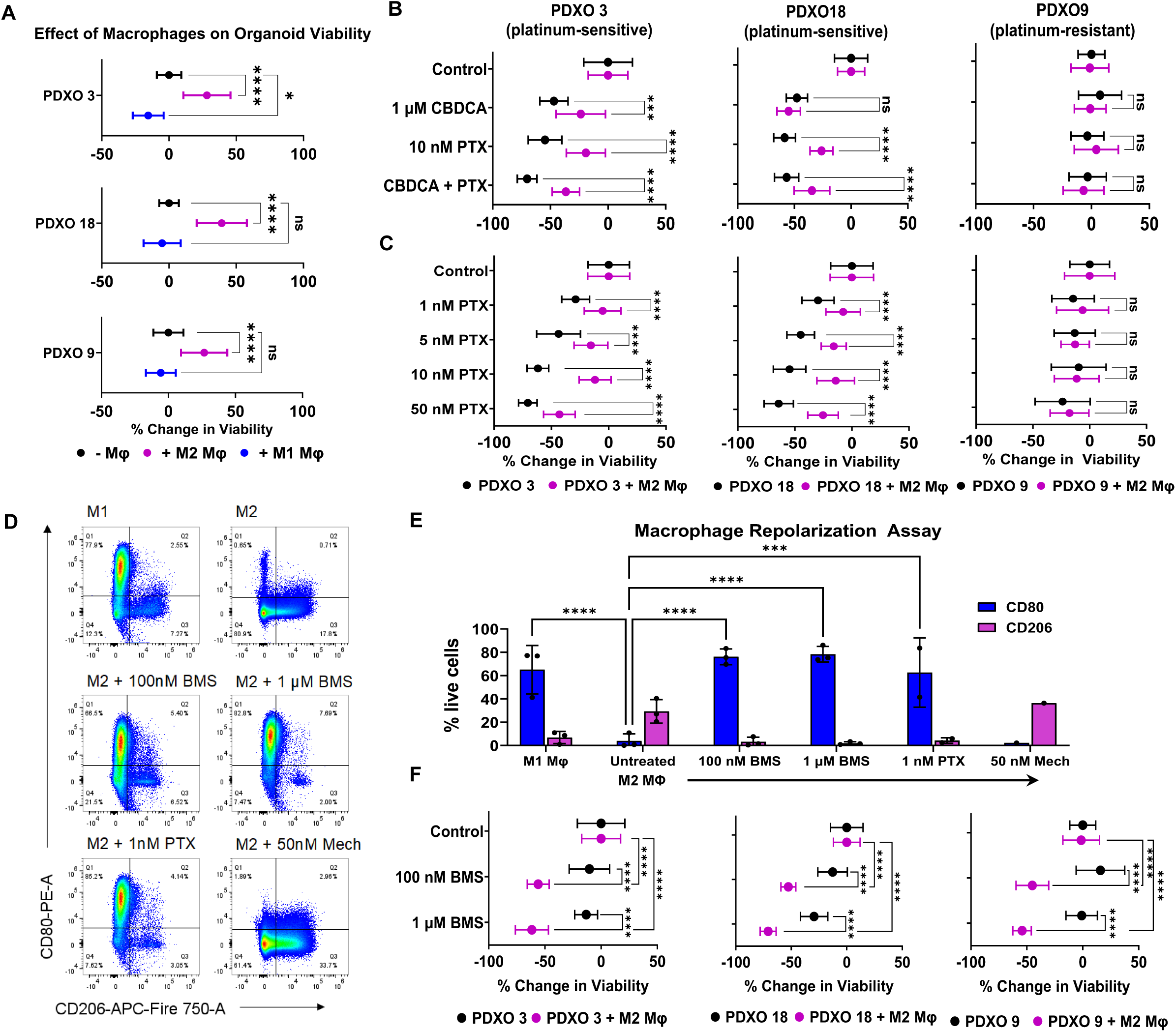
Impact of Macrophage Polarization on Response to Treatment in PDXO Models. **A.** Viability of PDXOs derived from solid tumors of platinum-sensitive or platinum-resistant patients co-cultured with or without THP-1-derived M1 or M2 macrophages, untreated. **B.** Change in cell viability for PDXOs co-cultured with or without THP-1-derived M2 macrophages, treated with 1 µM CBDCA, 10 nM PTX, or their combination, assessed over 72 hours. **C.** Viability of the same PDXOs treated with varying concentrations of PTX under similar co-culture conditions. **D.** THP-1-derived M2 macrophages were treated with BMS777607, PTX or mechlorethamine for 72 hours followed by spectral flow cytometry. Untreated M1 and M2 macrophages included as controls. Representative density plots show polarization states, measured by expression of CD80 (M1 marker) in Q1 and CD206 (M2 marker) in Q3. **E**. Average percentage of macrophages expressing CD80 (blue) and CD206 (purple), indicating repolarization. Data represent mean values from 1-3 experiments with standard deviations. Statistical significance is indicated by asterisks (ns: not significant; * p<0.05; ** p<0.01; *** p<0.001; **** p<0.0001), analyzed by two-way ANOVA. **F.** PDXOs treated with different concentrations of BMS777607, showing changes in viability normalized to untreated controls. For panels **A, B**, **C**, and **F**, data combines results from two independent experiments and at least 6 wells were tested/condition (technical replicates) in each experiment. The difference in viability is calculated relative to untreated controls (set to 0%), with more negative values indicating a stronger response to treatment. Error bars represent standard deviation. Statistical significance is indicated by asterisks (ns: not significant; * p<0.05; ** p<0.01; *** p<0.001; **** p<0.0001), analyzed by one-way ANOVA.

Organoids alone or co-cultured with M2 macrophages were treated with standard chemotherapeutic agents (carboplatin (CBDCA), paclitaxel (PTX), or their combination) for 72 hours and cell viability was assessed using the CellTiter 3D Glo assay. For equivalent comparison of treatment responses between organoids alone and organoids co-cultured with M2 macrophages, data were normalized to the untreated control (0% change) for each group.

As expected, platinum-sensitive organoid models alone, PDXO3 and PDXO18, responded to treatment with carboplatin, paclitaxel and the combination therapy, with a 50% or more reduction in viability (**Fig 3B**). Likewise, platinum-resistant PDXO9 showed no response, signifying agreement between the patient’s clinical designation and *in vitro* organoid response. PDXO3 and PDXO18 co-cultured with M2 macrophages showed improved viability when treated with paclitaxel alone or the combination therapy (**Fig 3B**). This effect was not observed in the platinum-resistant PDXO9, which is already resistant to both drugs. Co-culture with M2 macrophages had little effect on response to carboplatin, although viability was slightly higher in PDXO3 organoids co-cultured with M2 macrophages.

These findings highlight the influence of M2 macrophages on treatment efficacy. Combined with our data on differential PDX-specific recruitment of M2 macrophages, this underscores the need to consider the specific tumor environment when evaluating chemotherapy strategies, as the impact of immune cells like macrophages can significantly alter treatment outcomes.

### Impact of Macrophage-Targeted Therapies on Organoid Viability

It has been proposed that repolarizing TAMs from an M2-like phenotype to an M1-like phenotype should abrogate the effects of TAMs on OC proliferation and drug resistance, and previous studies have reported that low dose paclitaxel itself can repolarize TAMs (29). Therefore, to assess whether the effect of M2 macrophages on paclitaxel sensitivity is dependent on paclitaxel concentration, PDXOs with and without M2 macrophages were incubated in varying concentrations of paclitaxel (1 nM, 5 nM, 10 nM, 50 nM) for 72 hours (**Fig 3C**). For platinum-sensitive organoids, the presence of M2 macrophages increased cell viability regardless of paclitaxel concentration, although the greatest change in viability occurred at 10 nM paclitaxel.

We next tested whether other candidate therapies might be more effective at repolarizing macrophages. The tyrosine kinase receptor RON regulates macrophage polarization, skewing them towards an M2 phenotype (30). Therefore, we hypothesized that RON inhibition might more directly repolarize macrophages towards an M1-like, anti-tumor phenotype and abrogate macrophage-dependent paclitaxel resistance. The small molecule inhibitor, BMS777607, which inhibits RON and the related MET receptor as well as other immunosuppressive tyrosine kinase inhibitors, Tyro3, Axl, and Mertk, was chosen for testing. BMS777607 has been shown to improve response to a PD-1 inhibitor in a syngeneic mouse model of triple negative breast cancer (31). We first examined the ability of BMS777607 to repolarize THP-1-derived M2 macrophages. Other agents previously reported to have an immunomodulatory effect on macrophages, low dose paclitaxel (29, 32) and cyclophosphamide (33, 34) were included for comparison. Since cyclophosphamide is inactive *in vitro*, we used mechlorethamine, an analog of the active form of cyclophosphamide. THP-1 derived M2 macrophages were treated with 100 nM or 1 µM BMS777607, 1 nM low dose paclitaxel, or 50 nM mechlorethamine. After 72 hours, macrophages were harvested and the presence of CD80 (M1 marker) and CD206 (M2 marker) was analyzed by flow cytometry (**Fig 3D**). The percent of CD80+ and CD206+ cells were quantified for each samples (**Fig 3E**). Untreated THP-1 derived macrophages polarized to M1 or M2 were included as controls. Both concentrations of BMS777607 and 1 nM paclitaxel effectively shifted the polarization of M2 macrophages towards an M1 phenotype, as evidenced by increased CD80+ and decreased CD206+ macrophages. Mechlorethamine did not repolarize M2 macrophages.

Since we did not see a decrease in organoid viability with 1 nM paclitaxel treatment in co-culture experiments (**Fig 3C**), it is possible that the presence of cancer organoids counteracts the effects of paclitaxel-dependent macrophage repolarization. To determine whether BMS777607 can be effective in the presence of OC cells, PDXO organoids were cultured with or without M2 macrophages and then treated with BMS777607 for 72 hours. In the presence of M2 macrophages, organoids treated with BMS777607 showed a significant reduction in organoid viability in all three PDXOs, dropping viability to below 50% of the untreated controls (**Fig 3F**). Importantly, this reduction in organoid viability was seen in the platinum-resistant PDXO9 as well as in the platinum-sensitive PDXOs. BMS777607 treatment had no significant effect on the viability of organoids grown without macrophages, demonstrating that the effect of BMS777607 is targeted to TAMs. This study shows that BMS777607 has a potent effect on macrophages that in turn reduces organoid viability through limiting the release of pro-tumor growth factors and/or by stimulating an anti-tumor immune response.

### Immune Cell Distribution in OC Patient Ascites Fluid and Humanized PDX Models

To further explore TME dynamics and evaluate the impact of human TAMs on chemotherapy response, we moved to an *in vivo* humanized PDX model (huPDX3-Luc2). Human cord blood-derived CD34+ hematopoietic stem cells were engrafted into NBSGW immunocompromised mice and then PDX3-Luc2 OC spheroids were delivered by IP injection into the peritoneal cavity to model disseminated OC. These humanized PDX models have demonstrated recruitment of human immune cells to the TME and thus provide a relevant framework for studying human immune responses within the TME (25). Spectral flow analysis of the immune cell populations in the malignant ascites of OC patients at the time of surgery (**Fig 4A**) and huPDX models at endstage (**Fig 4B**) showed a similar distribution of myeloid cells, T cells, and B cells using a 14-color immune panel (Antibodies listed in **Table S1**). The immune cells from patient ascites samples were comprised of a high percentage of CD11b+ myeloid cells (30%-43%), CD8+ T cells (18%-30%), and CD4+ T cells (16%-23%), with low percentages of CD19+ B cells (**Fig 4A**). Likewise, human immune cells in the ascites of the three huPDX models were CD11b+ myeloid cells (39%-53%), CD8+ T cells (10%-18%) and CD4+ T cells (24%-52%) (**Fig 4B**).

**Figure 4.**
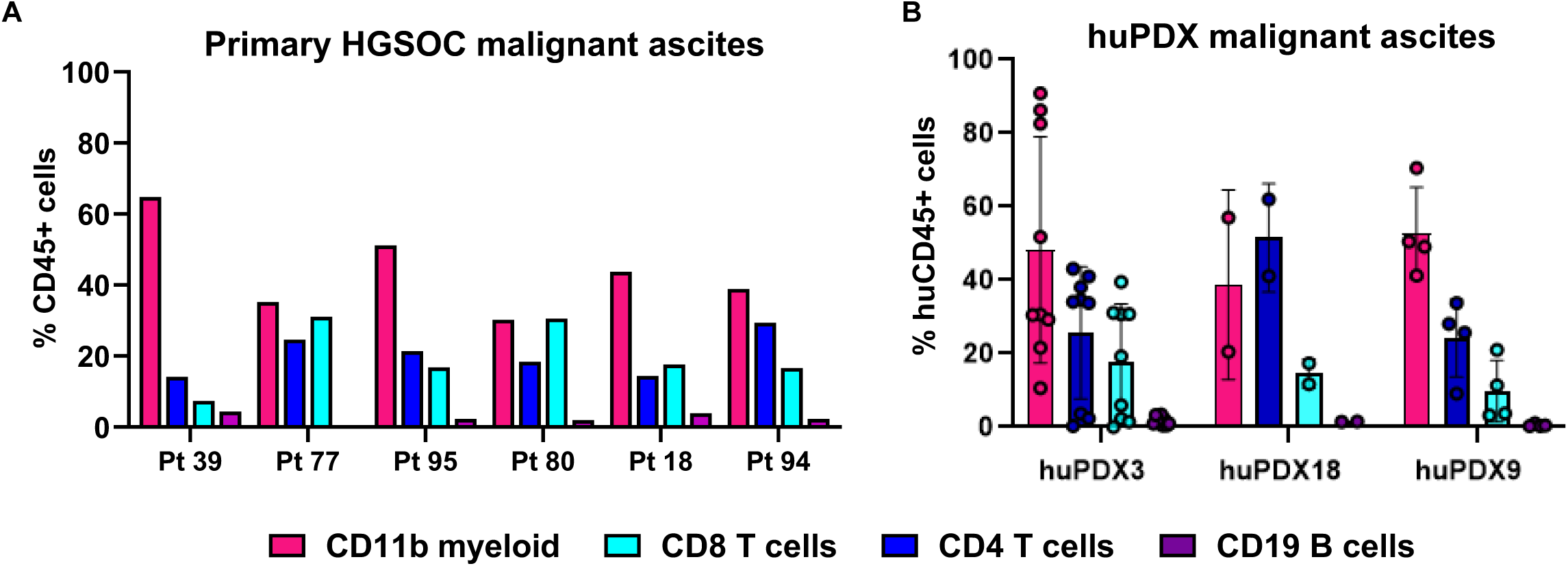
Immune Cell Distribution in OC Patient Ascites Fluid and Humanized PDX Models. **A.** Spectral flow analysis of CD45+ immune cells in the malignant ascites fluid of OC patients at time of surgery shows a high percentage of CD11b+ myeloid cells, CD4+ and CD8+ T cells and a low percentage of CD19+ B cells. **B**. Spectral flow analysis of human CD45+ cells in the malignant ascites of humanized PDX mice at end stage show a similar distribution to those of OC patients. Error bars are the standard deviation among mice in each group.

Importantly, this is a very different distribution of immune cells than has been found in the commonly used ID8 syngeneic mouse model that has been reported to have 60% T cells, 30% B cells, and less than 1% macrophages (35, 36) or in the BR5-akt syngeneic mouse model where the majority of mouse immune cells are MDSCs with <10% CD8 or CD4 T cells (37). These findings affirm that huPDX models are crucial for replicating the patient-specific immune TME, highlighting their value in translational cancer research. Their close correlation with patient samples demonstrates their usefulness for assessing how immune components influence chemotherapy response in a clinically relevant setting.

### Differential Chemotherapy Response in Non-Humanized vs. Humanized PDX Models

To test whether the presence of human immune cells altered response to chemotherapy, we implanted a platinum-sensitive OC PDX (PDX3) into non-humanized or humanized NBSGW and monitored change in tumor burden with treatment. We chose to use the platinum-sensitive OC PDX, PDX3, because it had demonstrated robust recruitment of human TAMs and T cells to the TME in a previous study (25). For non-humanized (non-hu) PDX and humanized PDX (huPDX) models, treatments were begun two weeks after PDX3-Luc2 implantation and included 12 mg/kg carboplatin (CBDCA) twice weekly, low-dose 6 mg/kg paclitaxel (PTX) daily, or combination paclitaxel plus carboplatin. A schematic of the study is shown in **Fig 5A**. For the huPDX study, we also examined the effect of TAM depletion in two additional groups treated with the CSF1R inhibitor, sotuletinib or BLZ945, (160 mg/kg, 5 days a week) alone or in combination with low dose paclitaxel. Studies in mouse models have demonstrated that CSF1R inhibition can either deplete mouse TAMs (38, 39) or repolarize them towards an M1-like anti-tumor phenotype (40).

**Figure 5.**
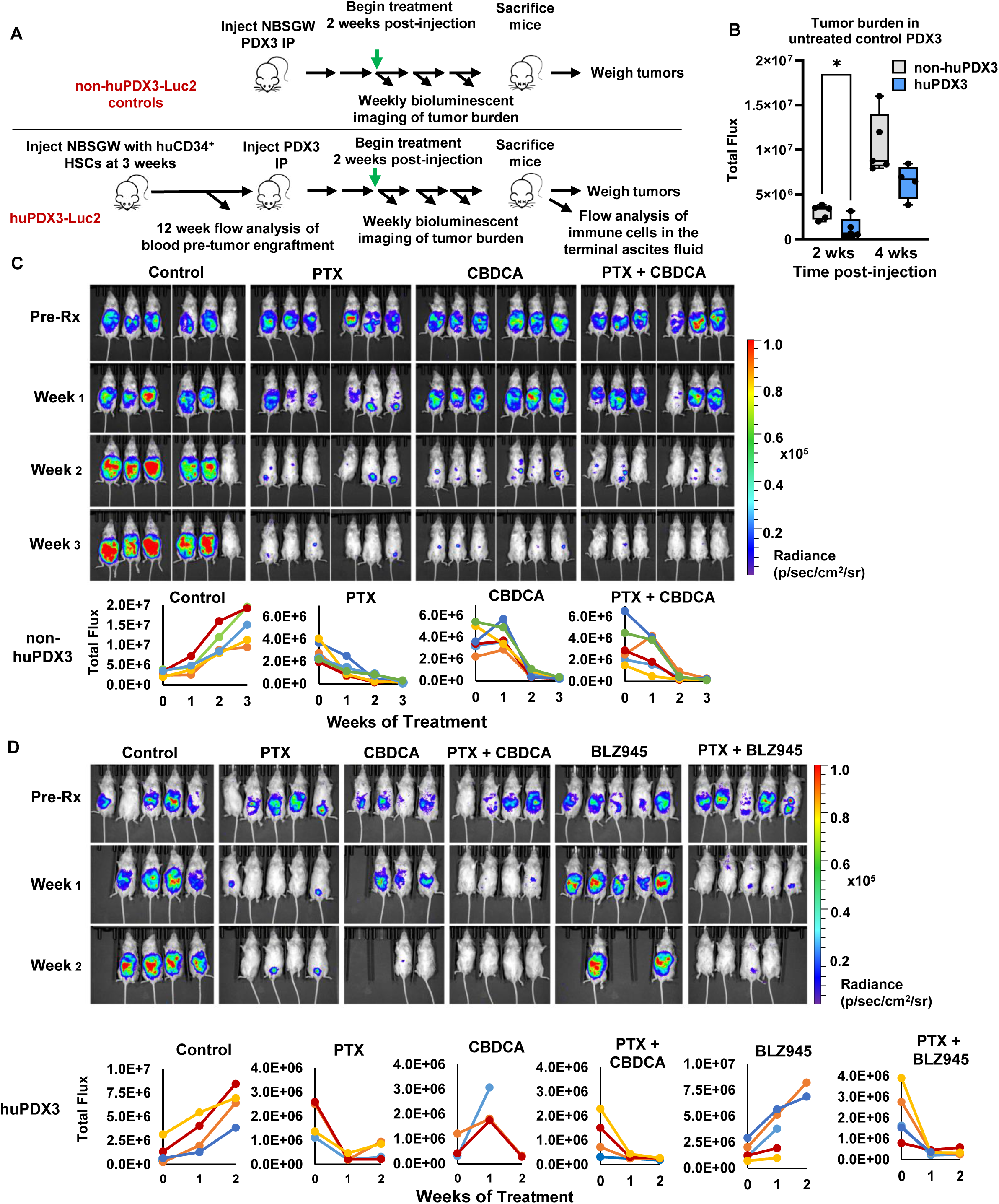
PDX3 Response to Standard Chemotherapy is Influenced by a Reconstituted Human Immune System. **A.** PDX3-Luc2 spheroids were injected into the peritoneal cavity of non-huPDX3 or huPDX3 NBSGW mice. Two weeks later, mice were treated with low-dose PTX, CBDCA or BLZ945, alone or combination. **B.** Tumor burden was quantified by bioluminescent imaging in control groups of non-huPDX3 and huPDX3 at 2 weeks vs 4 weeks post-engraftment. **C-D.** Bioluminescent imaging of tumor burden over time is shown for non-huPDX3 **(C)** and huPDX3 **(D)** mice. Graphs show **c**hange in tumor flux for each mouse/group.

Weekly bioluminescent imaging of tumor burden beginning pre-treatment allowed us to analyze treatment response in PDX with or without human immune cells. At pretreatment imaging, tumor burden was noticeably more variable among mice in huPDX3 compared to non-huPDX3. Although non-humanized and humanized mice were injected with the same number of PDX3-Luc2 cancer cells, untreated tumor burden at pre-treatment imaging (two weeks post-tumor implantation) was significantly less in huPDX compared to non-huPDX, suggesting that engrafted human immune cells initially suppress tumor growth (**Fig 5B**). However, by the 4^th^ week post-implantation, huPDX3 tumor burden in the untreated group was not significantly different than that of non-huPDX, supporting the idea that tumor cells manipulate the immune microenvironment, shifting human immune cells from anti-tumor to tumor-promoting (41).

Five huPDX3 mice across control, carboplatin, paclitaxel and paclitaxel plus BLZ945 groups had to be removed early from the study due to wasting and anemia. These are known symptoms that arise in humanized mice that develop a robust human macrophage population. Activated human macrophages attack mouse erythroid cells leading to anemia (42). Mice in the BLZ945 treatment group showed wasting, skin redness, and hair loss in patches, signs of immune activation that may be due to a systemic reduction in human immunosuppressive macrophages.

The presence of human immune cells in the huPDX3 mice affected the timing and degree of chemotherapy response. huPDX3 mice responded faster to paclitaxel and combination paclitaxel plus carboplatin treatment compared to non-huPDX3 (**Fig 5C, D**). After the first week of treatment, average tumor burden in huPDX3 treated with paclitaxel or the combination therapy was reduced to 17% +/- 12% and 24% +/- 8% respectively, compared to non-huPDX3 mice where average tumor burden was only reduced to 45% +/- 19% and 82% +/- 47%. However, by the second week, non-huPDX3 mice showed a reduction in tumor burden across chemotherapy treatment groups that continued into the third week of treatment.

Similar to our *in vitro* data, paclitaxel response appeared to be the most sensitive to the presence of human immune cells. After two weeks of treatment, two of the four huPDX3 mice in the paclitaxel treatment group exhibited an increase in tumor burden (tumor rebound)(**Fig 5D**). Paclitaxel response did not correlate with percent humanization since mice with 61% and 29% humanization in PBMCs at 16 weeks demonstrated tumor rebound, while mice with 82% and 42% humanization did not (data not shown). This rebound was dependent on human immune cells since the non-huPDX3 mice treated with paclitaxel showed a steady decrease in tumor burden over three weeks (**Fig 5C**). Overall, our findings highlight the significant role of human immune cells in modifying response to the standard chemotherapy, paclitaxel.

### Chemotherapy-Induced Modulation of Immune Cell Populations

At the experimental end point, malignant ascites fluid was collected and isolated cells underwent spectral flow analysis using a 15-color immune panel (**Table S1**) to examine changes in the peritoneal TME with treatment. Human CD45+ cells from all treatment groups were clustered and visualized by t-SNE (**Fig 6A**). Heatmaps of lineage marker expression on the t-SNE plot of all samples are shown in **Supplemental Fig 4**. Control huPDX ascites has three clusters of CD11b+ CD11c^high^ myeloid cells that are likely TAMs, as well as large clusters of CD4 and CD8 T cells and a small cluster of CD19+ B cells. Significant differences were observed with chemotherapy treatment compared to untreated controls, visualized in the t-SNE plots of each treatment group (**Fig 6B**). Quantification of population percentages are plotted by group (**Fig 6C, D**). huPDX ascites from carboplatin-treated mice had very few CD11b+ myeloid cells, while ascites from paclitaxel plus carboplatin-treated mice had significantly more myeloid cells compared to the control. For paclitaxel-treated huPDX, it was necessary to divide the group by response at two weeks, since the distribution of immune cells in the ascites was distinct in mice where tumor burden rebounded compared to mice with no rebound. Mice that rebounded had significantly more CD11b+ myeloid cells and fewer T cells, particularly CD4 T cells, than those that did not rebound. Macrophage marker expression was also different between the groups. Mice with no rebound had a significantly higher percentage of CD80+ and a lower percentage of CD86+ cells, both markers of anti-tumor TAMs. They also had a reduced percentage of CD206+ cells, but a higher percentage of CD163+ cells, markers of pro-tumor TAMs. Overall, paclitaxel treated mice that did not rebound had fewer myeloid cells and the remaining myeloid cells had an altered marker profile, though the change in markers did not clearly indicate an M2 to M1 repolarization as seen in our *in vitro* macrophage repolarization assay. A new cluster of CD11c^low^ myeloid cells was evident in ascites from any treatment group that included paclitaxel, suggesting that cells in this cluster are recruited or repolarized with paclitaxel treatment. However, there was no difference in the percentage of CD11c^low^ myeloid cells in mice with tumor rebound versus no rebound (**Fig 6D**).

**Figure 6.**
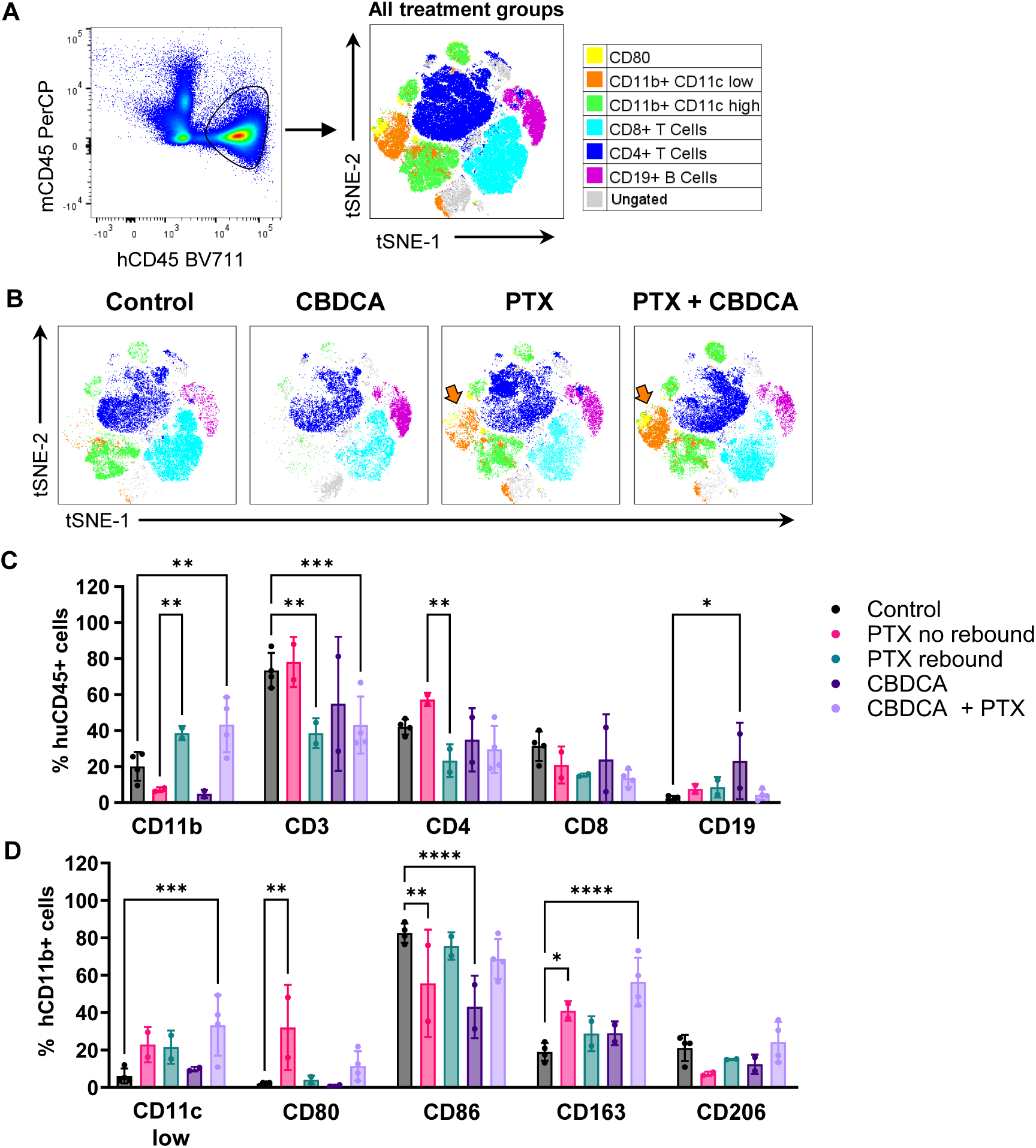
Chemotherapy-Induced Modulation of Immune Cell Populations. huPDX3 ascites fluid was analyzed by spectral flow cytometry. **A.** Representative dot plot of mouse CD45+ and human CD45+ cells from a single sample (left). Clusters of human CD45+ immune cells from all treatment groups (black circle) were visualized by t-SNE. Colors indicate cell subsets positive for the indicated immune cell marker. **B**. Clusters of immune cell populations from the ascites fluid of huPDX3 mice from the control and chemotherapy treatment groups: paclitaxel (PTX), carboplatin (CBDCA), and the combination (PTX + CBDCA). Colors are coded as in A. A cluster of CD11b+ CD11c low cells is specific to the groups with PTX treatment (orange arrow). **C.** Quantification of the percentage of hCD45+ myeloid cells (CD11b), T cells (CD3, CD4 and CD8) and B cells (CD19) for each treatment group. PTX treated mice were divided into mice with no rebound in tumor burden versus mice with rebound. **D.** CD11b+ myeloid cells were further classified with tumor associated macrophage (TAM) markers for CD11c, CD80 and CD86 (M1 markers), and CD163 and CD206 (M2 markers). Dots are the values for each mouse. Error bars are the SD among mice in each group. Statistical significance is indicated by asterisks (* p<0.05; ** p<0.01; *** p<0.001; **** p<0.0001), analyzed by one-way ANOVA.

These findings reinforce the idea that chemotherapy alters the immune microenvironment in a PDX model with human immune cells. It also raises the question of whether between-mouse variation in the immune environment may influence response to paclitaxel. While combination therapy also increased the percentage of myeloid cells, this group of huPDX did not show regrowth at two weeks of treatment. Our *in vitro* data indicated that PDX3 response to carboplatin therapy is less sensitive to the presence of macrophages (**Fig 3B**). Therefore, in the combination therapy, carboplatin may counteract the effect of human immune cells on paclitaxel response.

### Targeting Macrophage Recruitment as a Therapeutic Strategy

We have observed that macrophages promote paclitaxel resistance in PDX3 organoids. Therefore, we examined the effect of macrophage-targeted therapy on tumor growth and immune cell distribution in our huPDX study. Treatment with the CSF1R inhibitor BLZ945 alone did not reduce tumor burden (**Fig 5D**). Analysis of the immune cells in the ascites of BLZ945-treated mice showed that this therapy reduced immune cell numbers across all populations, not just macrophages. Of the two BLZ945-treated huPDX3 mice analyzed at end point, both had very few CD45+ cells in their ascites fluid (average 2.6% of total live cells), which is far lower than three of four control untreated huPDX (average 18.1%)(**Fig 7A**). t-SNE plot analysis (**Fig 7B**) and quantification of cell populations (**Fig 7C, D**) showed that the remaining immune cells in the BLZ945-treated ascites had a similar distribution to the untreated controls. BLZ945 appears to deplete macrophages and in the process limit the recruitment of other immune cells.

**Figure 7.**
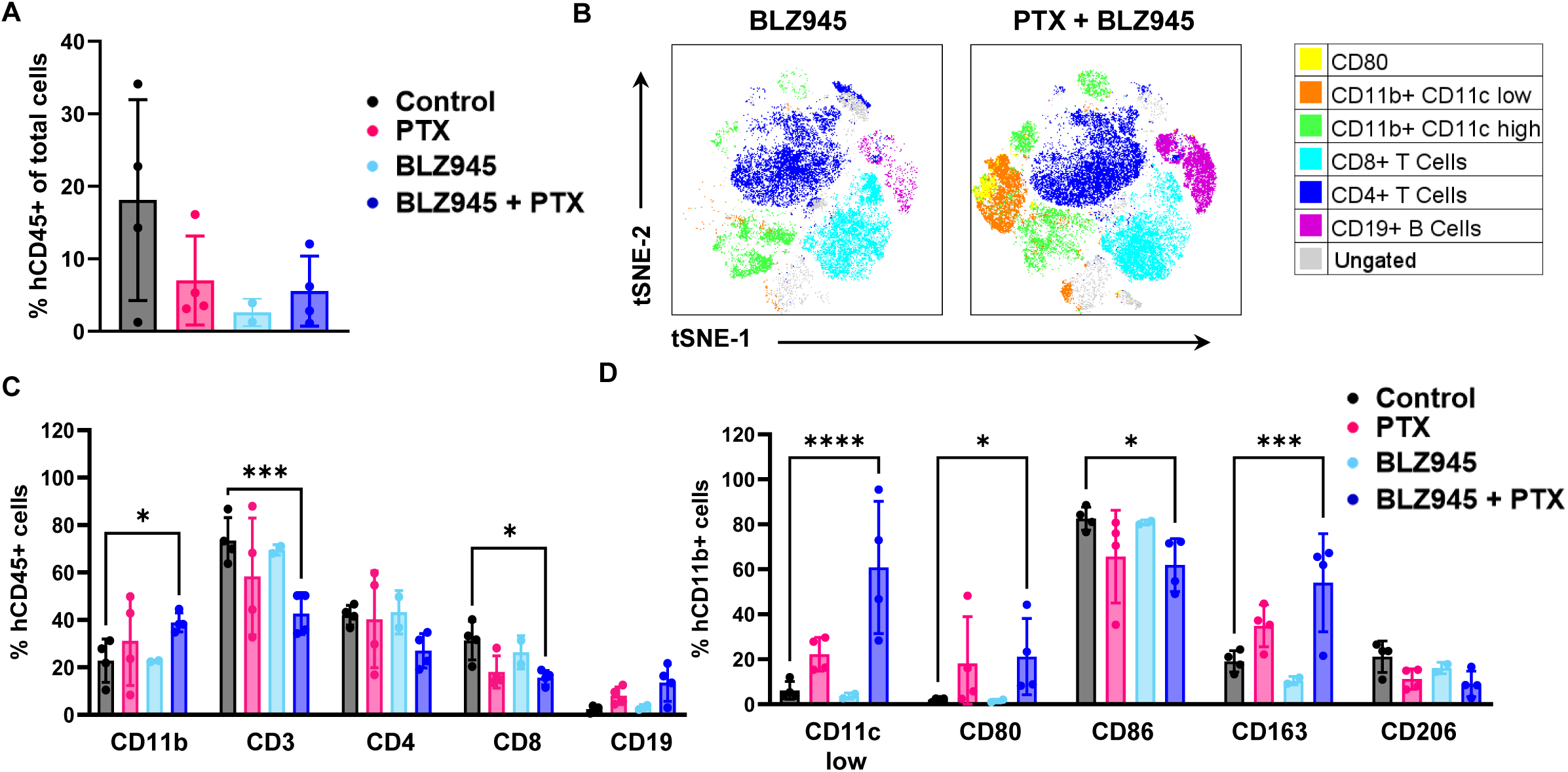
Targeting Macrophage Recruitment as a Therapeutic Strategy. Ascites fluid was collected from huPDX3 mice treated with low dose PTX, BLZ945, or BLZ945-PTX combination at endpoint and analyzed with spectral flow cytometry. **A.** The % hCD45+ cells per mouse in each treatment group compared to untreated control mice. **B.** t-SNE plots show clusters of immune cell populations in the ascites fluid of BLZ945 and BLZ945 + PTX treated mice. The PTX-specific cluster of CD11b+ CD11c low myeloid cells is present in mice from the double treatment group. **C.** Quantification of myeloid cells (CD11b), T cells (CD3, CD4, CD8) and B cells (CD19) for each treatment group. **D.** CD11b+ myeloid cells were further classified for low CD11c expression and for tumor associated macrophage (TAM) markers, CD80 and CD86 (M1 markers), and CD163 and CD206 (M2 markers). Dots are the values for each mouse. Error bars are the SD among mice in each group. Statistical significance is indicated by asterisks (* p<0.05; ** p<0.01; *** p<0.001; **** p<0.0001), analyzed by one-way ANOVA.

Tumors treated with the combination of BLZ945 and paclitaxel showed reduced tumor burden, similar to the paclitaxel monotherapy group (**Fig 5D**). Importantly, none of the mice in this group rebounded after two weeks of treatment, further supporting the idea that resistance to paclitaxel was macrophage-dependent. Ascites from this group had a distribution similar to paclitaxel and carboplatin combination therapy with a higher percentage of myeloid cells particularly in the CD11b+ CD11c^low^ cluster associated with paclitaxel treatment, a high percentage of CD80+ and CD163+ myeloid cell populations, and a lower percentage of T cells compared to untreated controls. These findings indicate that while macrophage-targeted BLZ945 might have a limited impact on PDX3 tumor burden as a monotherapy, combining BLZ945 with paclitaxel limits paclitaxel resistance and alters the ascites immune distribution.

In conclusion, our *in vitro* and *in vivo* studies reveal that M2 macrophages are preferentially recruited by cancer cells as shown in migration assays, and foster a tumor-permissive microenvironment that promotes resistance paclitaxel. We have identified the receptor tyrosine kinase inhibitor, BMS777607, as a possible macrophage repolarizing agent that effectively reduces organoid viability in the presence of M2 macrophages. Finally, our *in vivo* study comparing chemotherapy response in non-humanized versus humanized PDX models demonstrates the effects of human immune cells on chemotherapy response and vice versa, the effects of chemotherapy treatment on the immune cell distribution in OC ascites. These studies underscore the potential of immune modulation in combination with standard chemotherapy, as a promising strategy to enhance treatment efficacy.

## Discussion

Our previous work in huPDX models demonstrated that cancer cells from three OC PDX produced a distinct set of human cytokines that influenced immune cell recruitment in humanized mice (25). Here, using a 3D macrophage migration assay, we not only confirmed the findings from a recent study that OC cells are capable of recruiting macrophages within a 3D matrix (28), but went further to demonstrate that a significantly higher number of M2 macrophages are recruited to OC organoids compared to PBMC-derived monocytes or M1 macrophages. Higher levels of M2 macrophage recruitment were consistent across all five PDXO models and OVCAR8 organoids. M-CSF likely exerts a strong influence on macrophage recruitment, since the extent of M2 macrophage migration correlated with the M-CSF levels detected in the patient’s ascites fluid. This is also in agreement with an initial comparison of macrophage recruitment in two OC lines, with high versus low M-CSF levels (28). Surprisingly, higher levels of CCL2/MCP-1, a chemokine that recruits CCR2+ monocytes, did not improve monocyte migration in our assay. Our findings support the idea that M-CSF-secreting OC cancer cells preferentially recruit pro-tumor tissue-resident macrophages, perhaps from the peritoneal cavity, rather than recruiting circulating monocytes and polarizing them towards an M2-like phenotype as has been the accepted paradigm (43–47). However, our results do not rule out the possibility that some monocytes are recruited *in vivo* in the context of a full complement of immune cells. The principle role of OC-intrinsic M-CSF in recruiting M2 macrophages to the TME bears further study.

Interactions between OC cells and TAMs have been shown to play a significant role in treatment failure leading to relapse (48, 49). Our finding that co-culture with M2 macrophages reduces sensitivity to paclitaxel in PDXOs further supports the role of TAMs in paclitaxel resistance. We are the first to show that this effect is amplified in 3D spheroids and embedded organoids that more accurately represent the TME. We detected TAM-driven paclitaxel resistance in the well-studied OVCAR8 ovarian cancer cell line and in two paclitaxel-sensitive OC PDXOs. TAMs improve OC proliferation/survival across all models, and counteract the effects of paclitaxel in paclitaxel-sensitive OC.

Therapies that repolarize TAMs have the potential to transform the immunosuppressive TME into one that promotes an immune response. Our studies suggest that, while low dose paclitaxel and the tyrosine kinase inhibitor BMS777607 are both capable of repolarizing M2 macrophages *in vitro*, only BMS777607 was able to reduce the viability of PDXOs co-cultured with M2 macrophages. This reduction in viability occurred across all three PDXOs, including PDXO9 that demonstrated inherent carboplatin and paclitaxel resistance. Therefore, BMS777607 could be a potent and effective TAM-targeted therapy for OC treatment. Since BMS777607 is expected to repolarize rather than deplete TAMs, it may be more effective at promoting anti-tumor immune responses than BLZ945 that depletes rather than repolarizes macrophages. In a syngeneic mouse model of triple negative breast cancer, BMS777607 reduced tumor growth and improved response to PD-1 inhibition (31). Our *in vitro* data suggests that these combination immunotherapies may be effective in OC.

Our huPDX bridge the gap between traditional transgenic models and clinical disease by replicating the immune microenvironment observed in OC patients, as evidenced by the similar immune cell distribution seen in patient and huPDX ascites samples. Our *in vivo* studies used the OC huPDX3, a model that has demonstrated robust recruitment of human TAMs and T cells to the TME (27), allowing us to examine the interconnection of tumor associated immune cell recruitment and chemotherapy response. Here we showed that human immune cells increase the chance of paclitaxel resistance in the huPDX3 model. Tumor regrowth correlated with a change in the immune cell distribution in malignant ascites fluid characterized by an increase in TAMs and a reduction in T cells.

Our study is the first to examine the effects of CSF1R inhibition alone or in combination with paclitaxel in a huPDX mouse model. Previous studies in mouse models of glioblastoma and mammary tumors showed that treatment with CSF1R inhibitors depleted TAMs within the TME (38, 39). In our huPDX study, BLZ945 depleted TAMs, but also greatly reduced the total population of immune cells in the ascites fluid, highlighting the importance of TAMs in recruiting other immune cells to the OC TME. BLZ945 monotherapy did not reduce huPDX3 tumor burden, similar to what has been seen in a clinical trial in solid tumors (NCT02829723), where BLZ945 showed limited efficacy as a monotherapy or combined with a PD-1 inhibitor, PDR001. It is possible that BLZ945, by indirectly depleting tumor-infiltrating T cells, limits the effectiveness of a T cell-targeted PD-1 inhibitor.

Our findings that combination paclitaxel and BLZ945 treatment led to a consistent reduction in tumor burden are in agreement with previous studies in mouse mammary tumor models where BLZ945 improved paclitaxel response (19, 50). It was surprising that the combination therapy did not reduce the number of immune cells in the ascites fluid as much as with BLZ945 monotherapy. The new cluster of CD11b+ CD11c^low^ TAMs found in paclitaxel-treated mice may be less sensitive to CSF1R inhibition and thus are not depleted with BLZ945 treatment. Our data suggest that BLZ945 therapy may be more effective in combination therapies. Paclitaxel monotherapy is still occasionally used as second-line therapy for platinum-resistant OC. Our study suggests that using a macrophage-targeted therapy in combination with paclitaxel as an alternative to paclitaxel monotherapy, might improve response in these patients by limiting TAM-dependent paclitaxel resistance.

Our *in vivo* studies are proof of principle that the huPDX model can reveal important information about treatment response in the presence of human immune cells. However, the effect of chemotherapy treatment on tumor infiltrating immune cells likely varies among patients. Larger huPDX studies that compare response across a cohort of humanized PDX can begin to identify the mechanisms that lead to between-patient variability in response and could determine the extent to which paclitaxel plus macrophage-targeted therapies could be effective to limit acquired resistance and relapse in OC patients.

## Acknowledgments

We acknowledge support from the American Cancer Society IRG Pilot Grant (Steinkamp), the UNMCCC RAC Grant (Steinkamp), the UNM Pathology Department and the UNM Comprehensive Cancer Center (UNMCCC), and a UNMCCC Pilot Grant to support funding for P. Nikeghbal. We would like to acknowledge the UNMCCC Support Grant NCI P30CA118100 that supports the UNMCCC Shared Resources. We would like to particularly thank Irina Lagutina PhD and Lillian Fitzpatrick in the Animal Models Shared Resource, Michael Paffett PhD in the Fluorescene Microscopy Shared Resource, Wade M Johnson PhD, Victoria Balise PhD, and Yuanyuan Gao PhD in the Flow Shared Resource. We would also like to thank Kristina Thiel PhD (University of Iowa) and Kim Leslie MD (UNM) for support with organoid development and Shayna Lucero and Rachel Grattan for technical support.

