## Supplemental Figures for "The Influence of Macrophages within the Tumor Microenvironment on Ovarian Cancer Growth and Response to Therapies"

### Supplemental Table 1. Antibodies Used for Flow Cytometry.

| Marker-fluorochrome | Catalog number | Clone | Dilution |
|---|---|---|---|
| huCD163-BV421 | 333612 | GHI/61 | 1:200 |
| huCD3-BV605 | 300460 | UCHT1 | 1:330 |
| huCD4-BV650 | 300536 | RPA-T4 | 1:1,000 |
| huCD45-BV711 | 304049 | HI30 | 1:200 |
| huPD-1-BV750 | 329965 | EH12.2H7 | 1:500 |
| huCD11c-AF488 | 301617 | 3.9 | 1:1,000 |
| muCD45-PerCP | 103129 | 30-F11 | 1:2,000 |
| huCD66b-PerCP-Cy5.5 | 305108 | G10F5 | 1:500 |
| hCD80-PE | 305207 | 2D10 | 1:500 |
| huCD19-PE Dazzle594 | 302251 | HIB19 | 1:1,000 |
| huCD11b-PE-Cy7 | 301322 | ICRF44 | 1:500 |
| huCD8-APC | 344721 | SK1 | 1:500 |
| hCD206-APC-Fire 750 | 321133 | 15-2 | 1:500 |
| huCD86-647 | 376304 | QA19A61 | 1:250 |

**Supplemental Table 1. Antibodies Used for Flow Cytometry.** 13 human-specific markers targeting immune cell surface proteins and a viability stain (all antibodies from BioLegend, San Diego, CA)

**Supplemental Figure 1. Polarization of Macrophages into M1 and M2.**

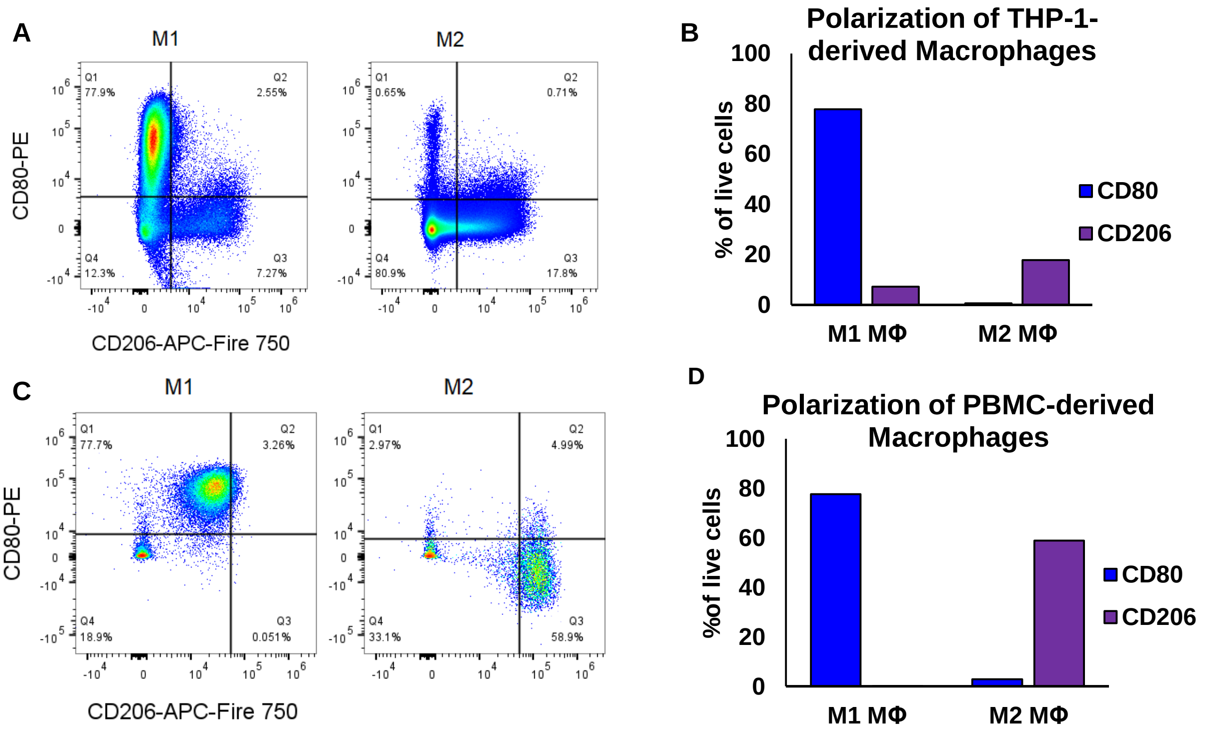


**Supplemental Figure 1. Polarization of Macrophages into M1 and M2.** Macrophages were derived from both THP-1 cells **(A-B)** and PBMCs **(C-D)** and differentiated into either M1 or M2 cells. **A and C.** Live cells from M1 and M2 macrophages analyzed with spectral flow cytometry for CD80 (M1 marker) and CD206 (M2 marker) to check for polarization. **B and D.** Quantification of polarized cells from one experiment.

**Supplemental Figure 2. Equal Distribution of Humanization Across all HuPDX3 Groups.**

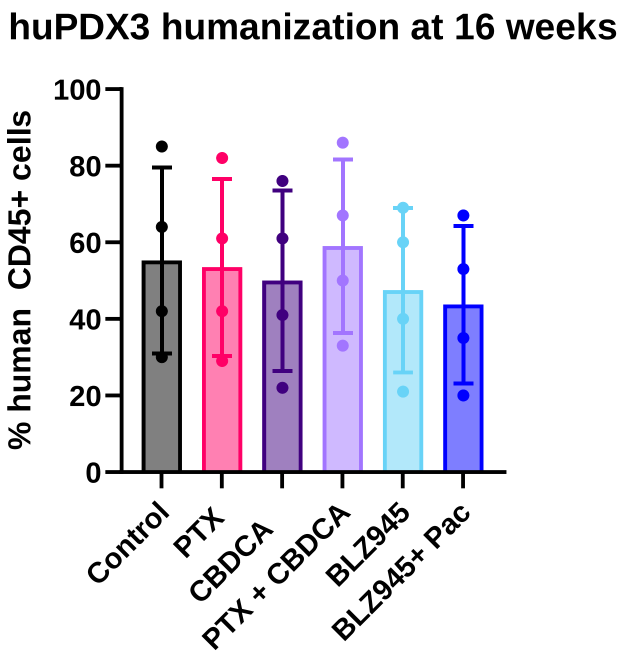


**Supplemental Figure 2. Equal Distribution of Humanization Across all HuPDX3 Groups.** Blood was drawn at 16 weeks and analyzed by flow using human CD45 and mouse CD45 fluorescently tagged antibodies. Humanization of CD45 cells in peripheral blood was determined by dividing the number of human CD45+ cells by the sum of the human and mouse CD45+ cells.

**Supplemental Figure 3. The Effect of Macrophages on Viability at Start of Experiment.**

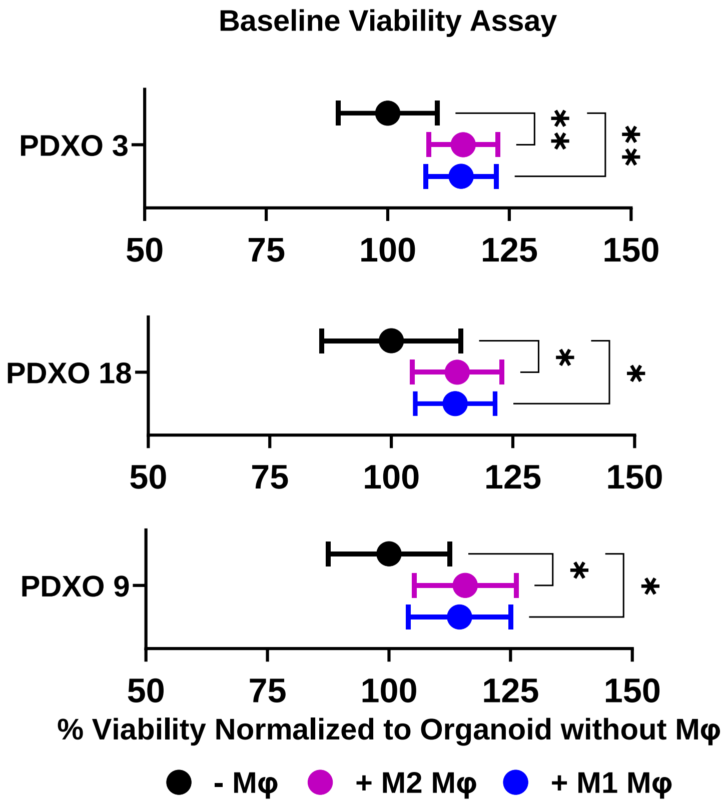


**Supplemental Figure 3. The Effect of Macrophages on Viability at Start of Experiment.** Whole-well viability measurement of PDXOs seeded with or without THP-1-derived M1 or M2 macrophages and assessed on Day 0. The increase in viability between PDXO – macrophages and + macrophages should represent the metabolic activity of the macrophages.

**Supplemental Figure 4. Expression of Lineage Markers to Identify Myeloid, T Cell and B Cell Populations in HuPDX3 Ascites.**

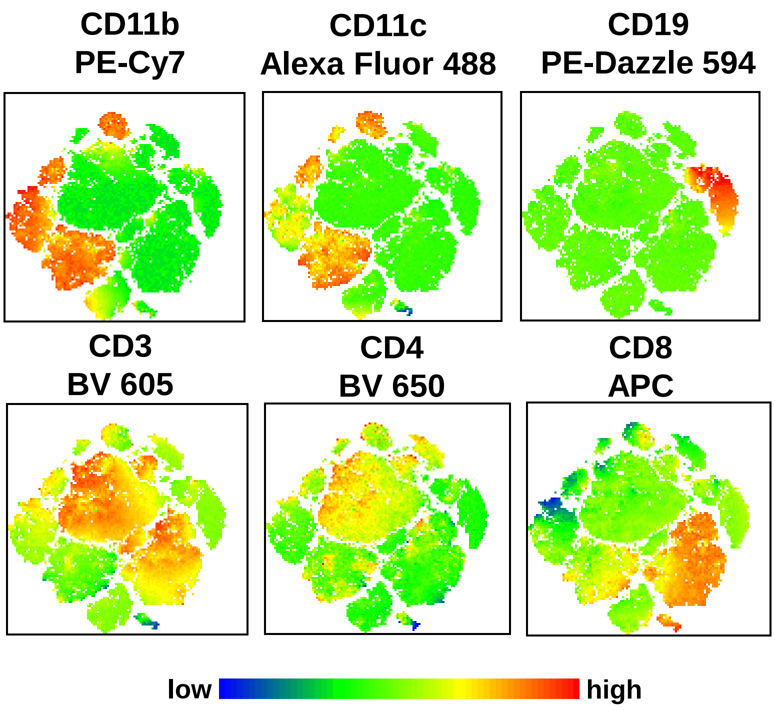


**Supplemental Figure 4. Expression of Lineage Markers to Identify Myeloid, T Cell and B Cell Populations in HuPDX3 Ascites.** CD45+ cells from all huPDX3 ascites samples (control, PTX, CBDCA, BLZ945, CBDCA + PTX, BLZ945 + PTX) were concatenated into a single file (n=20) and visualized by t-SNE. Heatmaps of lineage markers were used to identify myeloid, T cell and B cell clusters.
